## Supplementary Information for "Wakefulness Induced by TAAR1 Partial Agonism is Mediated Through Dopaminergic Neurotransmission"

**This file includes:**

Figures S1 to S4
Tables S1 to S2

### Supplementary Figures

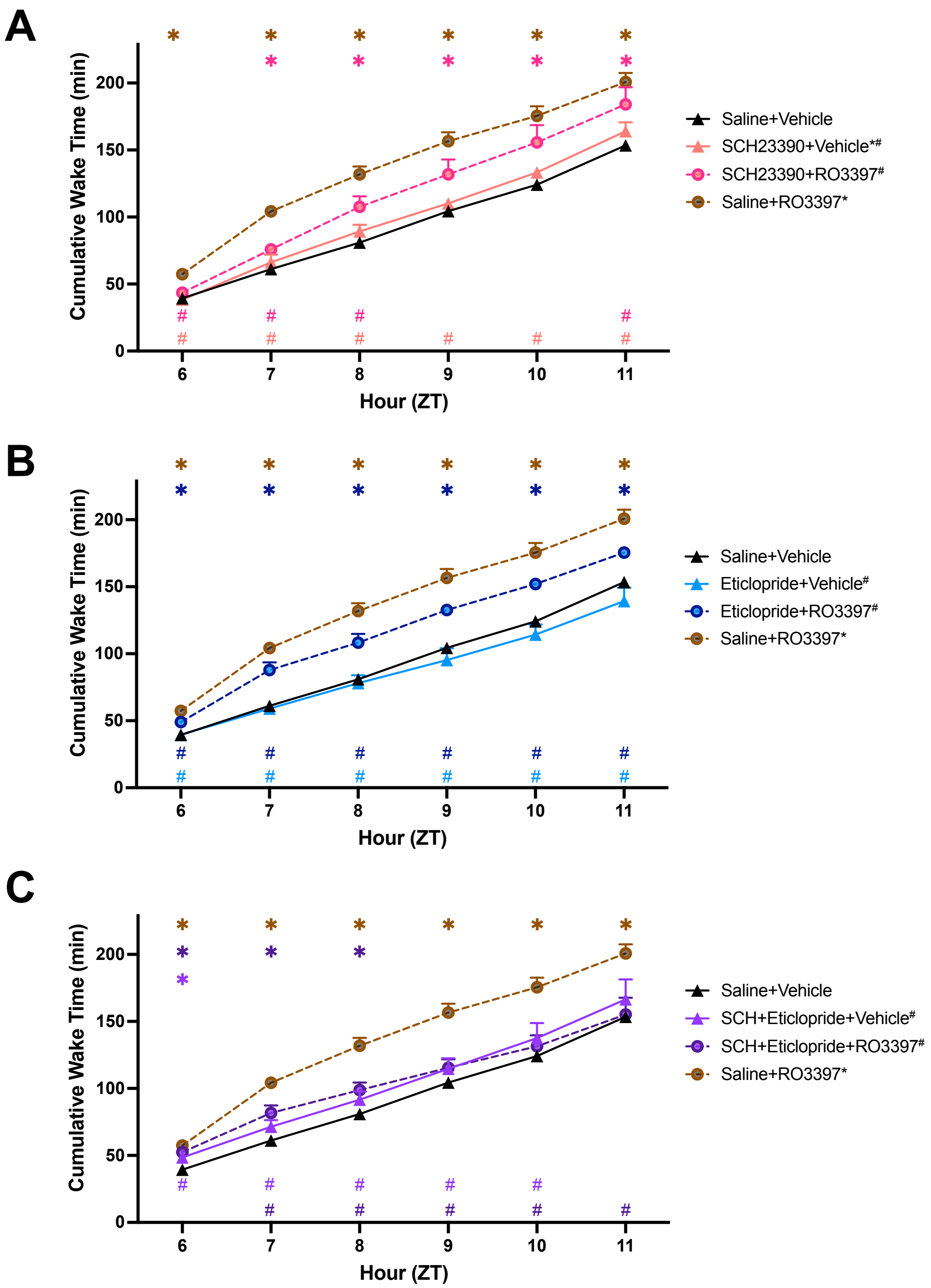


**Figure S1. Cumulative Wake time for the first 6 hours after the second dosing.** For ease of visualization, data are split into three subgroups in which the results from the negative (Sal+Veh) and positive (Sal+RO3397) control treatments are repeated in each graph. **A-C**. Cumulative Wake time (mean+SEM). Colored symbols indicate statistical significance for that hour compared to Sal+Veh(*) or Sal+RO3397(#) based on RM-ANOVA. *, # $p < 0.05$;

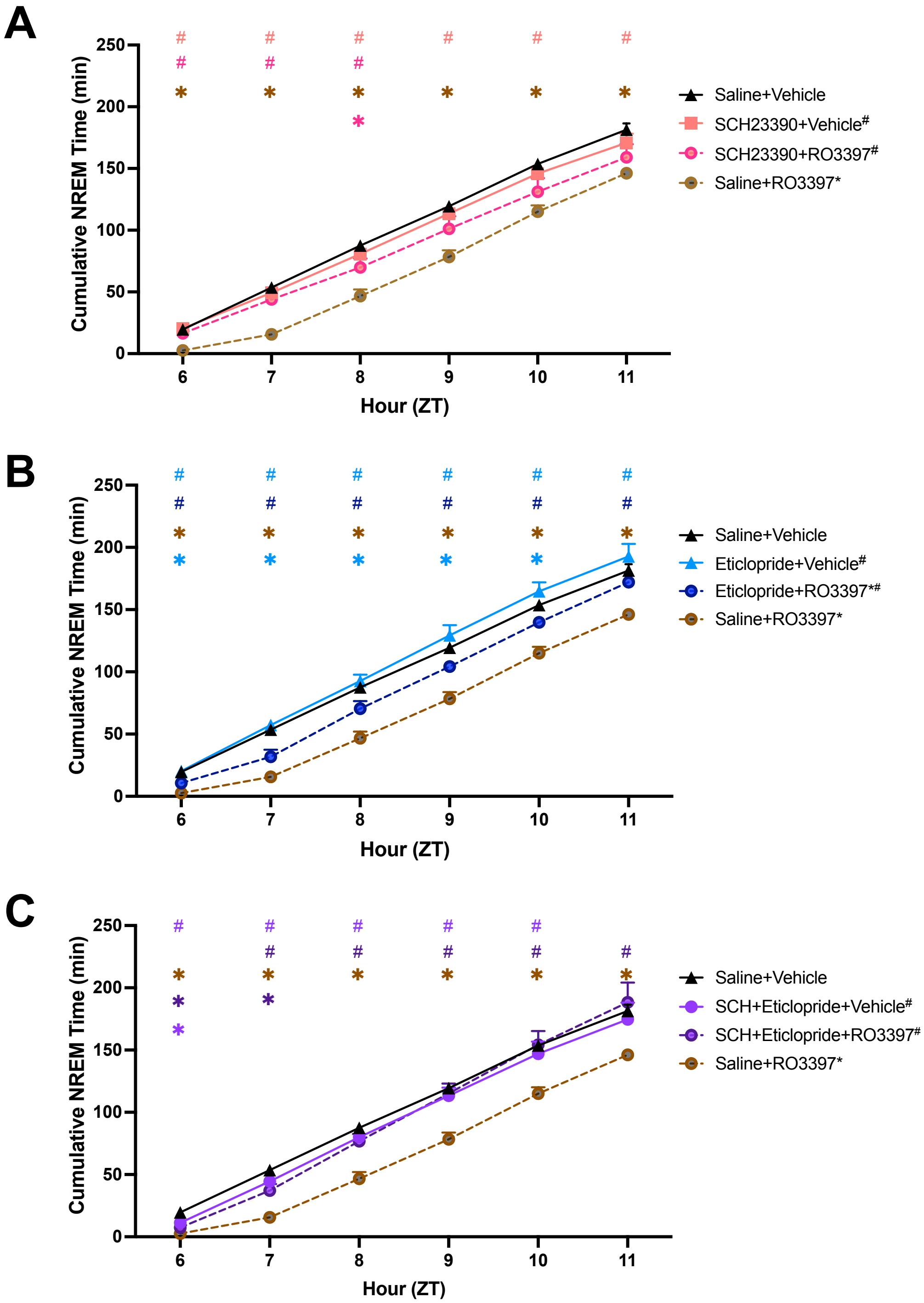


**Figure S2. Cumulative NREM time for the first 6 hours after the second dosing.** For ease of visualization, data are split into three subgroups in which the results from the negative (Sal+Veh) and positive (Sal+RO3397) control treatments are repeated in each graph. **A-C**. Cumulative NREM time (mean+SEM). Colored symbols indicate statistical significance for that hour compared to Sal+Veh(*) or Sal+RO3397($^{\#}$) based on RM-ANOVA. *, $^{\#}$ $p < 0.05$;

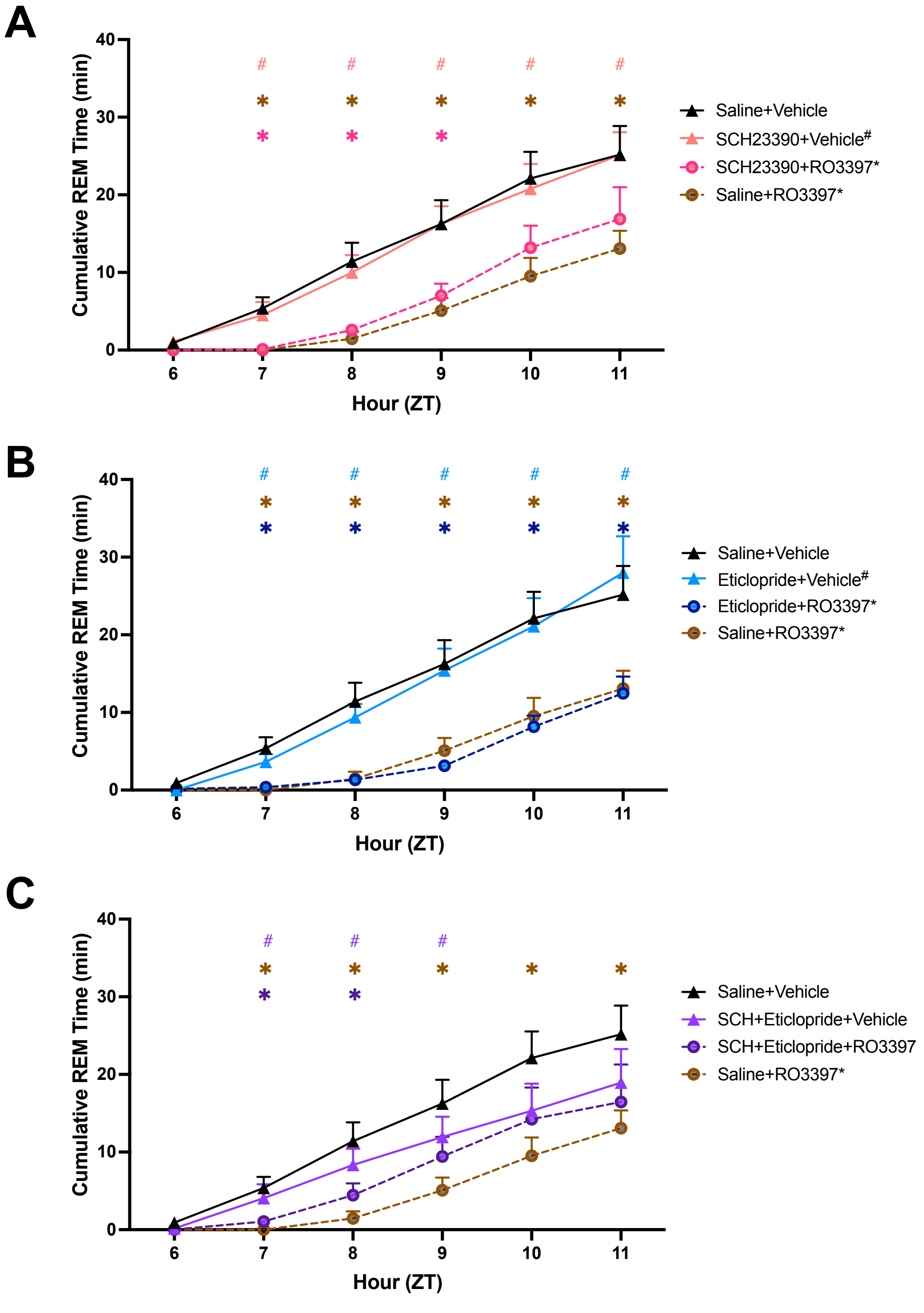


**Figure S3. Cumulative REM time for the first 6 hours after the second dosing.** For ease of visualization, data are split into three subgroups in which the results from the negative (Sal+Veh) and positive (Sal+RO3397) control treatments are repeated in each graph. **A-C**. Cumulative REM time (mean+SEM). Colored symbols indicate statistical significance for that hour compared to Sal+Veh(*) or Sal+RO3397(#) based on RM-ANOVA. *, # $p < 0.05$;

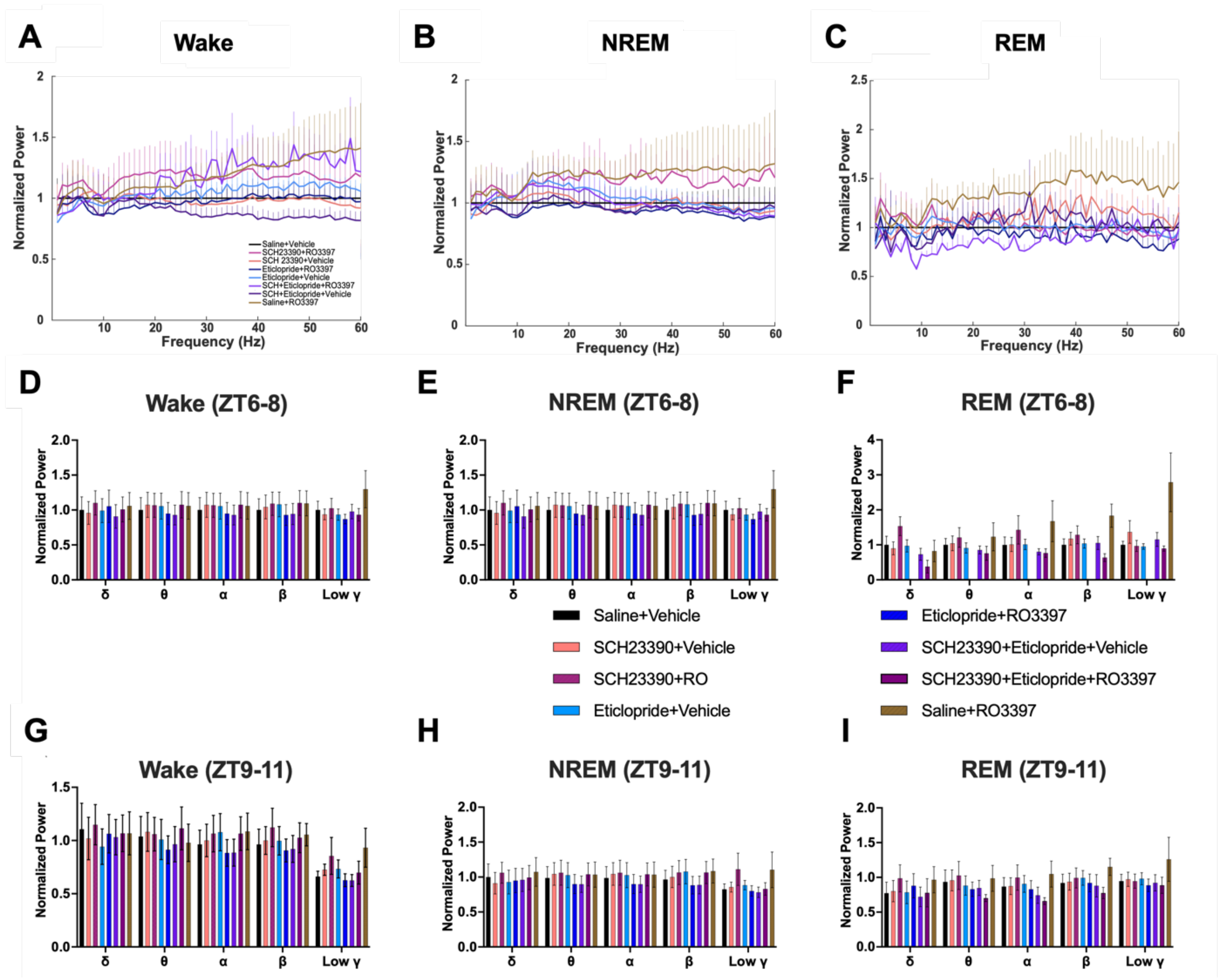


**Figure S4. EEG spectral power (0-60 Hz) for each treatment during the first 6 post-dosing hours. A-C.** 6 hour average spectral power for (A) Wake, (B) NREM sleep and (C) REM sleep. **D-F**. Normalized EEG power binned into the conventional bandwidths for each treatment during the first 3 post-dosing hours (ZT6-8) for (D) Wake, (E) NREM sleep and (F) REM sleep. **G-I**. Normalized EEG power binned into the conventional bandwidths for each treatment during the second 3 post-dosing hours (ZT9-11) for (G) Wake (H) NREM sleep and (I) REM sleep.

**Table S1.** Statistics underlying Figures and Supplementary Figures.

| Figure/Panel (Time bin) | Data Structure | Type of test | *F* value | *p* value | Variable |
|---|---|---|---|---|---|
| **1B-D** | normal distribution | 2-way ANOVA (Condition) | 3.919 | 0.0019 | ZT6-ZT11 hourly wake time (min) |
| 1B | normal distribution | Tukey's multiple comparison | | <0.0001 | Saline+Vehicle vs. Saline+RO3397 |
| 1C | normal distribution | Tukey's multiple comparison | | 0.0060 | Eticlopride+RO3397 vs. Saline+RO3397 |
| 1D | normal distribution | Tukey's multiple comparison | | 0.0075 | SCH23390+eticlopride+RO3397 vs. Saline+RO3397 |
| **1E-1G** | normal distribution | 2-way ANOVA (Condition x Time) | 5.235 | 0.0002 | ZT6-8. ZT9-11 3-hour bin total wake time (min) |
| 1E,1F,1G | normal distribution | Tukey's multiple comparison | | <0.0001 | Saline+Vehicle vs. Saline+RO3397 |
| 1E | normal distribution | Tukey's multiple comparison | | 0.0002 | SCH23390+Vehicle vs. Saline+RO3397 |
| 1F | normal distribution | Tukey's multiple comparison | | <0.0001 | Eticlopride+Vehicle vs. Saline+RO3397 |
| 1G | normal distribution | Tukey's multiple comparison | | 0.0005 | SCH23390+eticlopride+Vehicle vs. Saline+RO3397 |
| 1G | normal distribution | Tukey's multiple comparison | | 0.0086 | SCH23390+eticlopride+RO3397 vs. Saline+RO3397 |
| **2A-C** | heterogeneous dist | Brown-Forsythe ANOVA test | 15.990 | <0.0001 | ZT6-12 NREM latency |
| | | Welch's ANOVA | 12.850 | <0.001 | ZT6-12 NREM latency |
| 2A,B,C | normal distribution | Dunnett's multiple comparison | | 0.0005 | Saline+Vehicle vs. Saline+RO3397 |
| 2A | normal distribution | Dunnett's multiple comparison | | 0.0003 | SCH23390+Vehicle vs. Saline+RO3397 |
| 2A | normal distribution | Dunnett's multiple comparison | | 0.0004 | SCH23389+RO3397 vs. Saline+RO3397 |
| 2B | normal distribution | Dunnett's multiple comparison | | 0.0003 | Eticlopride+Vehicle vs. Saline+RO3397 |
| 2C | normal distribution | Dunnett's multiple comparison | | 0.0076 | SCH23390+eticlopride+Vehicle vs. Saline+RO3397 |
| **2D-F** | normal distribution | 2-way ANOVA (Condition) | 2.391 | 0.0350 | ZT6-NT12 hourly NREM time (min) |
| 2D,E,F | normal distribution | Tukey's multiple comparison | | 0.0003 | Saline+Vehicle vs. Saline+RO3397 |
| 2D | normal distribution | Tukey's multiple comparison | | 0.0164 | SCH23390+Vehicle vs. Saline+RO3397 |
| 2E | normal distribution | Tukey's multiple comparison | | 0.0013 | Eticlopride+Vehicle vs. Saline+RO3397 |
| 2E | normal distribution | Tukey's multiple comparison | | 0.0023 | Eticlopride+RO3397 vs. Saline+RO3397 |
| 2F | normal distribution | Tukey's multiple comparison | | 0.0256 | SCH23390+eticlopride+RO3397 vs. Saline+RO3397 |
| **2G-I** | normal distribution | 2-way ANOVA (Condition x Time) | 4.179 | 0.0012 | ZT6-9, ZT9-12 3-hour bin total NREM time (min) |
| 2G,2H,2I | normal distribution | Tukey's multiple comparison | | 0.0004 | Saline+Vehicle vs. Saline+RO3397 |
| 2G | normal distribution | Tukey's multiple comparison | | 0.0060 | SCH23390+Vehicle vs. Saline+RO3397 |
| 2H | normal distribution | Tukey's multiple comparison | | <0.0001 | Eticlopride+Vehicle vs. Saline+RO3397 |
| 2I | normal distribution | Tukey's multiple comparison | | 0.0078 | SCH23390+eticlopride+Vehicle vs. Saline+RO3397 |
| 2I | normal distribution | Tukey's multiple comparison | | 0.0246 | SCH23390+eticlopride+RO3397 vs. Saline+RO3397 |
| **3A-C** | heterogeneous dist | Brown-Forsythe ANOVA test | 10.550 | <0.0001 | ZT6-12 REM latency |
| | | Welch's ANOVA | 8.901 | <0.0001 | ZT6-12 REM latency |
| 3A,B,C | normal distribution | Dunnett's multiple comparison | | 0.0019 | Saline+Vehicle vs. Saline+RO3397 |
| 3A | normal distribution | Dunnett's multiple comparison | | 0.0013 | SCH23390+Vehicle vs. Saline+RO3397 |
| 3B | normal distribution | Dunnett's multiple comparison | | 0.0106 | Eticlopride+Vehicle vs. Saline+RO3397 |
| 3C | normal distribution | Dunnett's multiple comparison | | 0.0466 | SCH23390+eticlopride+Vehicle vs. Saline+Vehicle |
| 3C | normal distribution | Dunnett's multiple comparison | | 0.0233 | SCH23390+eticlopride+Vehicle vs. Saline+RO3397 |
| **3D-F** | normal distribution | 2-way ANOVA (Condition x Time) | 2.458 | 0.0307 | NT6-NT12 NREM hourly time (min) |
| 3D,E,F | normal distribution | Tukey's multiple comparison | | 0.0167 | Saline+Vehicle vs. Saline+RO3397 |
| 3D | normal distribution | Tukey's multiple comparison | | 0.0073 | SCH23390+Vehicle vs. Saline+RO3397 |
| 3E | normal distribution | Tukey's multiple comparison | | 0.0146 | Eticlopride+Vehicle vs. Saline+RO3397 |
| **3G-I** | normal distribution | 2-way ANOVA (Condition x Time) | 2.473 | 0.0299 | ZT6-9, ZT9-12 3-hour bin total NREM time (min) |
| 3G,H,I | normal distribution | Tukey's multiple comparison | | 0.0470 | Saline+Vehicle vs. Saline+RO3397 |
| **4A-C** | normal distribution | 2-way ANOVA (Condition x Time) | 2.490 | 0.0290 | ZT6-ZT12 Raw Temperature |
| 4B | normal distribution | Tukey's multiple comparison | | 0.0170 | Eticlopride+Vehicle vs. Saline+Vehicle |
| 4C | normal distribution | Tukey's multiple comparison | | 0.0130 | SCH23390+eticlopride+Vehicle vs. Saline+RO3397 |
| 4C | normal distribution | Tukey's multiple comparison | | 0.0320 | SCH23390+eticlopride+RO3397 vs. Saline+RO3397 |
| **4D-F** | normal distribution | 2-way ANOVA (Condition x Time) | 9.660 | <0.0001 | ZT6-ZT12 LMA (counts/min) |
| 4D,E,F | normal distribution | Tukey's multiple comparison | | 0.0467 | Saline+Vehicle vs. Saline+RO3397 |
| 4E | normal distribution | Tukey's multiple comparison | | 0.0001 | Eticlopride+Vehicle vs. Saline+Vehicle |
| 4E | normal distribution | Tukey's multiple comparison | | 0.0004 | Eticlopride+RO3397 vs. Saline+Vheicle |
| 4E | normal distribution | Tukey's multiple comparison | | 0.0014 | Eticlopride+Vehicle vs. Saline+RO3397 |
| 4E | normal distribution | Tukey's multiple comparison | | 0.0061 | Eticlopride+RO3397 vs. Saline+RO3397 |
| 4F | normal distribution | Tukey's multiple comparison | | 0.0002 | SCH23390+eticlopride+Vehicle vs. Saline+Vehicle |
| 4F | normal distribution | Tukey's multiple comparison | | 0.0001 | SCH23390+eticlopride+RO3397 vs. Saline+Vehicle |

| Figure/Panel (Time bin) | Data Structure | Type of test | *F* value | *p* value | Variable |
|---|---|---|---|---|---|
| 4F | normal distribution | Tukey's multiple comparison | | 0.0001 | SCH23390+eticlopride+RO3397 vs. Saline+Vehicle |
| 4F | normal distribution | Tukey's multiple comparison | | 0.0030 | SCH23390+eticlopride+Vehicle vs. Saline+RO3397 |
| 4F | normal distribution | Tukey's multiple comparison | | 0.0013 | SCH23390+eticlopride+RO3397 vs. Saline+RO3397 |
| **Fig.S1** | normal distribution | 2-way ANOVA (Condition x Time) | 2.882 | <0.0001 | ZT6-11 Cumulative Wake Time |
| Fig.S1A,B,C | normal distribution | Tukey's multiple comparison | | <0.0001 | Saline+Vehicle vs. Saline+RO3397 |
| Fig.S1A | normal distribution | Tukey's multiple comparison | | <0.0001 | SCH23390+Vehicle vs. Saline+RO3397 |
| Fig.S1A | normal distribution | Tukey's multiple comparison | | 0.0464 | SCH23390+RO3397 vs. Saline+RO3397 |
| Fig.S1B | normal distribution | Tukey's multiple comparison | | <0.0001 | Eticopride+Vehicle vs. Saline+RO3397 |
| Fig.S1B | normal distribution | Tukey's multiple comparison | | 0.0033 | Eticlopride+RO3397 vs. Saline+RO3397 |
| Fig.S1C | normal distribution | Tukey's multiple comparison | | 0.0025 | SCH23390+eticlopride+Vehicle vs. Saline+RO3397 |
| Fig.S1C | normal distribution | Tukey's multiple comparison | | 0.0016 | SCH23390+eticlopride+RO3397 vs. Saline+RO3397 |
| **Fig.S2** | normal distribution | 2-way ANOVA (Condition x Time) | 5.474 | 0.0010 | ZT6-11 Cumulative NREM Time |
| Fig.S2A | normal distribution | 2-way ANOVA (Condition x Time) | | 0.0001 | SCH23390+Vehicle vs. Saline+RO3397 |
| Fig.S2A | normal distribution | Tukey's multiple comparison | | 0.0401 | SCH23390+RO3397 vs. Saline+RO3397 |
| Fig.S2B | normal distribution | Tukey's multiple comparison | | 0.0001 | Eticopride+Vehicle vs. Saline+RO3397 |
| Fig.S2B | normal distribution | Tukey's multiple comparison | | 0.0007 | Eticlopride+RO3397 vs. Saline+RO3397 |
| Fig.S2C | normal distribution | Tukey's multiple comparison | | 0.0025 | SCH23390+eticlopride+Vehicle vs. Saline+RO3397 |
| Fig.S2C | normal distribution | Tukey's multiple comparison | | 0.0050 | SCH23390+eticlopride+RO3397 vs. Saline+RO3397 |
| 3H (ZT19-ZT24) | normal distribution | 2-way ANOVA (Condition x Time) | 4.761 | <0.0001 | ZT19-ZT24 Normalized NR Beta Power |
| 3H (ZT1-ZT6) | normal distribution | 2-way ANOVA (Condition x Time) | 2.543 | 0.0008 | ZT1-ZT6 Normalized NR Beta Power |
| 3H (ZT7-ZT12) | normal distribution | 2-way ANOVA (Condition x Time) | 1.589 | 0.0632 | ZT7-ZT12 Normalized NR Beta Power |
| 4A (ZT19-ZT24) | normal distribution | 2-way ANOVA (Condition x Time) | 2.660 | 0.0027 | WT mice Cumulative W Time (min) |
| 4B (ZT19-ZT24) | normal distribution | 2-way ANOVA (Condition x Time) | 2.780 | 0.0003 | HET mice Cumulative W Time (min) |
| 4C (ZT19-ZT24) | normal distribution | 2-way ANOVA (Condition x Time) | 1.040 | 0.4274 | KO mice Cumulative W Time (min) |
| 4D (ZT19-ZT24) | normal distribution | 2-way ANOVA (Condition x Time) | 1.980 | 0.0266 | WT mice Cumulative NR Time (min) |
| 4E (ZT19-ZT24) | normal distribution | 2-way ANOVA (Condition x Time) | 2.870 | 0.0002 | HET mice Cumulative NR Time (min) |
| 4F (ZT19-ZT24) | normal distribution | 2-way ANOVA (Condition x Time) | 1.010 | 0.4548 | KO mice Cumulative NR Time (min) |
| 4G (ZT19-ZT24) | normal distribution | 2-way ANOVA (Condition x Time) | 1.510 | 0.1211 | WT mice Cumulative REM Time (min) |
| 4H (ZT19-ZT24) | normal distribution | 2-way ANOVA (Condition x Time) | 1.820 | 0.0253 | HET mice Cumulative REM Time (min) |
| 4I (ZT19-ZT24) | normal distribution | 2-way ANOVA (Condition x Time) | 1.270 | 0.2173 | KO mice Cumulative REM Time (min) |

**Table S2.** Basal sleep/wake parameters of male C57BL6/J mice treated with a dopaminergic antagonist followed by TAAR1 agonist or vehicle.

| | | Time (min) | | Number of bouts | | Mean Bout Duration (min) | |
|---|---|---|---|---|---|---|---|
| Treatment Condition | N | ZT6-8 | ZT9-11 | ZT6-8 | ZT9-11 | ZT6-8 | ZT9-11 |
| ***WAKE*** | | | | | | | |
| Saline+Vehicle | 7 | 81.0 ± 3.4 | 72.5 ± 2.6 | 28.4 ± 3.0 | 29.0 ± 3.0 | 3.0 ± 0.5 | 2.6 ± 0.3 |
| SCH23390+Vehicle | 7 | 89.2 ± 5.0$^{\dagger\dagger\dagger}$ | 74.9 ± 5.6 | 36.0 ± 2.8 | 25.6 ± 1.5 | 2.4 ± 0.2 | 2.9 ± 0.3 |
| SCH23390+RO3397 | 7 | 107.6 ± 7.8 | 76.5 ± 7.2 | 37.0 ± 3.3 | 31.3 ± 4.8$^{\dagger\dagger\dagger\dagger}$ | 2.9 ± 0.3 | 2.6 ± 0.4 |
| Eticlopride+Vehicle | 7 | 78.0 ± 5.9$^{\dagger\dagger\dagger\dagger}$ | 61.3 ± 5.4 | 52.9 ± 4.2$^{**}$ | 46.7 ± 4.6$^{\dagger\dagger\dagger\dagger}$ | 1.4 ± 0.2$^{\dagger\dagger\dagger\dagger}$ | 1.3 ± 0.1 |
| Eticlopride+RO3397 | 7 | 108.3 ± 6.6 | 67.2 ± 7.6 | 56.4 ± 5.8$^{***}$ | 45.7 ± 5.1$^{\dagger\dagger\dagger\dagger}$ | 2.0 ± 0.3$^{\dagger\dagger\dagger\dagger}$ | 1.6 ± 0.4 |
| SCH23390+Eticlopride+Vehicle | 7 | 91.6 ± 6.1$^{\dagger\dagger\dagger}$ | 74.9 ± 10.4 | 57.3 ± 8.7$^{***}$ | 52.71 ± 4.14$^{\dagger\dagger\dagger\dagger}$ | 1.8 ± 0.3$^{\dagger\dagger\dagger\dagger}$ | 1.4 ± 0.2 |
| SCH23390+Eticlopride+RO3397 | 7 | 98.8 ± 5.7$^{\dagger\dagger}$ | 56.4 ± 8.4 | 57.0 ± 4.8$^{***}$ | 45.9 ± 6.3$^{\dagger\dagger\dagger\dagger}$ | 1.8 ± 0.2$^{\dagger\dagger\dagger\dagger}$ | 1.2 ± 0.1 |
| Saline+RO3397 | 7 | 131.9 ± 5.9$^{****}$ | 68.9 ± 3.1 | 33.0 ± 3.6$^{**}$ | 35.3 ± 5.7 | 4.3 ± 0.6 | 2.1 ± 0.3 |
| 2-way ANOVA | | $F_{(7,48)}$ = 3.919; p = 0.002 | | $F_{(7,48)}$ = 21.19; p < 0.0001 | | $F_{(7,48)}$ = 9.56; p < 0.0001 | |
| ***NREM Sleep*** | | | | | | | |
| Saline+Vehicle | 7 | 87.6 ± 2.5 | 93.8 ± 3.1 | 48.9 ± 4.6 | 49.0 ± 6.0 | 1.9 ± 0.3 | 2.1 ± 0.3 |
| SCH23390+Vehicle | 7 | 80.8 ± 3.8$^{\dagger\dagger}$ | 90.0 ± 6.1 | 53.1 ± 4.1 | 43.6 ± 3.1 | 1.6 ± 0.2 | 2.1 ± 0.1 |
| SCH23390+RO3397 | 7 | 69.8 ± 7.3 | 89.2 ± 4.6 | 47.6 ± 4.0 | 48.1 ± 5.3 | 1.5 ± 0.2 | 2.0 ± 0.3 |
| Eticlopride+Vehicle | 7 | 92.6 ± 5.2$^{\dagger\dagger\dagger\dagger}$ | 100.1± 6.4 | 72.7 ± 5.3$^{*\dagger\dagger\dagger\dagger}$ | 68.4 ± 3.8 | 1.3 ± 0.1 | 1.5 ± 0.1 |
| Eticlopride+RO3397 | 7 | 70.4 ± 6.1 | 101.7 ± 8.6 | 65.7 ± 3.9$^{\dagger\dagger\dagger}$ | 67.4 ± 5.2 | 1.0 ± 0.1$^{*}$ | 1.5 ± 0.1 |
| SCH23390+Eticlopride+Vehicle | 7 | 80.1 ± 5.0$^{\dagger\dagger}$ | 94.5 ± 10.2 | 71.9 ± 6.1$^{*\dagger\dagger\dagger\dagger}$ | 72.7 ± 6.0* | 1.1 ± 0.1 | 1.3 ± 0.2 |
| SCH23390+Eticlopride+RO3397 | 7 | 76.8 ± 5.9$^{\dagger}$ | 111.6 ± 11.02 | 61.7 ± 7.6$^{\dagger\dagger}$ | 68.4 ± 5.6 | 1.3 ± 0.2 | 1.7 ± 0.3 |
| Saline+RO3397 | 7 | 46.7 ± 5.3$^{***}$ | 99.5 ± 3.0 | 32.4 ± 3.3 | 55.7± 5.5 | 1.4 ± 0.1 | 1.9 ± 0.2 |
| 2-way ANOVA | | $F_{(7,48)}$ = 2.391; p = 0.035 | | $F_{(7,48)}$ = 8.347; p < 0.0001 | | $F_{(7,48)}$ = 2.50; p = 0.0283 | |
| ***REM Sleep*** | | | | | | | |
| Saline+Vehicle | 7 | 11.4 ± 2.4 | 13.8 ± 2.1 | 12.0 ± 3.3 | 15.1 ± 2.2 | 1.0 ± 0.2 | 0.8 ± 0.1 |
| SCH23390+Vehicle | 7 | 10.0± 2.2 | 15.1 ± 1.2 | 9.0 ± 2.1 | 14.6 ± 1.6 | 1.0 ± 0.1 | 1.0 ± 0.1 |
| SCH23390+RO3397 | 7 | 2.6 ± 0.7 | 14.3 ± 3.9 | 2.6 ± 0.7 | 14.1 ± 4.2 | 1.1 ± 0.3 | 1.0 ± 0.1 |
| Eticlopride+Vehicle | 7 | 9.4± 1.7 | 18.6 ± 3.2 | 11.4 ± 2.1 | 24.6 ± 4.5 | 0.8 ± 0.1 | 0.8 ± 0.1 |
| Eticlopride+RO3397 | 7 | 1.3 ± 0.5* | 11.1 ± 1.9 | 1.7 ± 0.7 | 17.9 ± 2.4 | 0.4 ± 0.0$^{*}$ | 0.5 ± 0.1 |
| SCH23390+Eticlopride+Vehicle | 7 | 8.4 ± 2.1 | 10.6 ± 3.2 | 15.4 ± 3.6 | 18.4 ± 5.9 | 0.4 ± 0.0$^{*}$ | 0.5 ± 0.1 |
| SCH23390+Eticlopride+RO3397 | 7 | 4.5 ± 1.5 | 12.0 ± 3.6 | 7.3 ± 2.6 | 16.7 ± 5.4 | 0.4 ± 0.0$^{*}$ | 0.6 ± 0.1 |
| Saline+RO3397 | 7 | 1.4 ± 0.9* | 11.6 ± 1.5 | 1.9 ± 0.8 | 12.9 ± 1.5 | 0.6 ± 0.2 | 0.86 ± 0.07 |
| 2-way ANOVA | | $F_{(7,48)}$ = 2.473; p = 0.030 | | $F_{(7,48)}$ = 2.204; p = 0.050 | | $F_{(7,48)}$ = 7.80; p < 0.0001 | |

*p < 0.05; **p < 0.01; ***p < 0.001; ****p < 0.0001 vs. Saline+Vehicle

$^{\dagger}p < 0.05$; $^{\dagger\dagger}p < 0.01$; $^{\dagger\dagger\dagger}p < 0.001$; $^{\dagger\dagger\dagger\dagger}p < 0.0001$ vs. Saline+RO3397
